# Automatic reconstruction of metabolic pathways from identified biosynthetic gene clusters

**DOI:** 10.1101/2020.11.24.395400

**Authors:** Snorre Sulheim, Fredrik A. Fossheim, Alexander Wentzel, Eivind Almaas

## Abstract

**Background:** A wide range of bioactive compounds are produced by enzymes and enzymatic complexes encoded in biosynthetic gene clusters (BGCs). These BGCs can be identified and functionally annotated based on their DNA sequence. Candidates for further research and development may be prioritized based on properties such as their functional annotation, (dis)similarity to known BGCs, and bioactivity assays. Production of the target compound in the native strain is often not achievable, rendering heterologous expression in an optimized host strain as a promising alternative. Genome-scale metabolic models are frequently used to guide strain development, but large-scale incorporation and testing of heterologous production of complex natural products in this framework is hampered by the amount of manual work required to translate annotated BGCs to metabolic pathways. To this end, we have developed a pipeline for an automated reconstruction of BGC associated metabolic pathways responsible for the synthesis of non-ribosomal peptides and polyketides, two of the dominant classes of bioactive compounds.

**Results:** The developed pipeline correctly predicts 72.8% of the metabolic reactions in a detailed evaluation of 8 different BGCs comprising 228 functional domains. By introducing the reconstructed pathways into a genome-scale metabolic model we demonstrate that this level of accuracy is sufficient to make reliable *in silico* predictions with respect to production rate and gene knockout targets. Furthermore, we apply the pipeline to a large BGC database and reconstruct 943 metabolic pathways. We identify 17 enzymatic reactions using high-throughput assessment of potential knockout targets for increasing the production of any of the associated compounds. However, the targets only provide a relative increase of up to 6% compared to wild-type production rates.

**Conclusions:** With this pipeline we pave the way for an extended use of genome-scale metabolic models in strain design of heterologous expression hosts. In this context, we identified generic knockout targets for the increased production of heterologous compounds. However, as the predicted increase is minor for any of the single-reaction knockout targets, these results indicate that more sophisticated strain-engineering strategies are necessary for the development of efficient BGC expression hosts.

## Background

Natural products provide an immense source of bioactive small molecules of medical and agricultural importance [1, 2, 3]. The biosynthesis of these small-molecule bioactive compounds is usually governed by genes that are clustered in physical close proximity on the genome in fungal [4] or bacterial species [5], commonly known as biosynthetic gene clusters (BGCs). The revolution in sequencing technology has enabled access to complete genome sequences for an increasing number of bacteria and fungi. Mining of these genomes has revealed a vast abundance of BGCs, many more than the number of bioactive compounds observed *in vitro* [6, 7], suggesting that many BGCs are not expressed or that their respective compounds are not produced at detectable amounts in laboratory conditions. The activation of these silent BGCs may lead to the discovery of many novel bio-pharmaceuticals [8].

One promising avenue towards exploration of the bioactive potential of these silent BGCs is heterologous expression in host strains that are engineered to achieve maximal production of the encoded natural products [9, 10]. With current software [11] it is possible to quickly mine a genome for BGCs and retrieve information about the class, location, and functional domains of every gene in each cluster [12]. One may further prioritize BGC candidates for heterologous expression based on this information, (dis)similarity to known BGCs, bioactivity assays and mass spectrometry profiles of produced compounds, and subsequently transfer the selected BGCs to a chosen host strain using available genetic tools [13, 14]. However, the cloning and transfer of BGCs can be time-consuming and difficult depending on the genetic tools available for the native and the heterologous host strains, as well as the size of the BGC in question [15]. Additionally, it is not clear which host strain or which genetic modifications will maximize the yield of the secondary metabolite synthesized through the metabolic pathway catalyzed by the enzymes, or enzyme complexes, encoded by the heterologously expressed BGC [16, 17].

Genome-scale metabolic models (GEMs) can predict the consequence of genetic modifications [18] and are routinely used to guide strain design for a wide range of purposes [19]. However, this approach has still not gained traction in guiding strain-engineering efforts to increase the heterologous production of complex natural compounds, despite a number of available GEMs for *Actinobacteria* [20], a phylum known for an extremely diverse secondary metabolism responsible for about two-thirds of all known antibiotics in use today [21]. Previous efforts are limited to maximization of native secondary metabolites [22, 23, 24] or precursor pools [25]. One reason for the lack of computational efforts leveraging GEMs to assess heterologous production from BGCs is the significant amount of work required to map out the associated metabolic pathway, although most of the required information is contained in the output from software used to identify and annotate BGCs, such as antiSMASH [12]. In this work, we address this hurdle by developing a pipeline that parses the output obtained from antiSMASH and constructs the corresponding metabolic-synthesis pathway, thereby making BGCs available for constraint-based analysis and strain engineering guided by GEMs.

We have chosen to focus on non-ribosomal peptide synthetases (NRPSs) and two types of polyketide synthases (PKSs), namely Type 1 PKSs and trans-AT PKSs. These BGC classes are of particular interest because of their vast abundance [26, 27] and great prospect to become novel biopharmaceuticals [28, 29]. For an exhaustive description of NRPS and PKS biosynthesis, we refer the reader to a range of excellent reviews [30, 31, 32, 27, 33], but we provide the brief summary required as a context for the later description of the pipeline and results. Both NRPS and PKS biosynthesis are performed by multidomain enzyme complexes that create a polymer from amino acid or acyl-CoA building blocks, respectively. The chain elongation is modular, and in general collinear, so that the length of the final polymer corresponds to the number of active elongating modules. This makes it tractable to predict the biosynthetic pathways producing the associated compounds from the annotated sequence data. An active chain elongating module in an NRPS cluster requires at least three functional domains: a condensation (C) domain, an adenylation domain and a peptidyl carrier (PCP) domain. The A domain activates a specific amino acid (or in some cases a carboxylic acid) and facilitates the attachment of the amino acid to the PCP domain, while the C domain catalyzes the formation of peptide bonds required to elongate the peptide. In addition to these three domains, NRPS modules can replace the C domain by a Cy domain performing condensation and heterocyclisation or additionally contain a methyltransferase (MT) and/or an epimerase (E) domain. The load module initiating biosynthesis usually lacks the C domain, while the terminating module contains either a thioesterase (TE) or a thioester reductase (TR) domain.

Similar to NRPSs, chain elongating modules of PKSs rely on three functional domains: an acyltransferase (AT) domain that recognizes a specific extender unit and attaches it to the acyl carrier (ACP) domain. The third domain, ketosynthase (KS) catalyzes the Claissen condensation required to extend the polyketide chain. A standard PKS load module contains only the AT and ACP domain, and a TE or TR domain is required for the release of the polyketide chain by the final PKS module. PKS modules can also feature the reducing domains ketoreductase (KR), dehydratase (DH) and enoylreductase (ER), and different combinations of functional domains yield a large variety of molecular transformations, in particular for the trans-AT PKSs [32]. These trans-AT PKSs not only differ from normal (*cis*) modular PKSs by having a larger module diversity and deviations from canonical rules, but they are also recognized by freestanding AT domains that perform the chain elongation [32]. The diversity of PKS and NRPS natural products is further extended by hybrid variants containing both NRPS and PKS domains and modules.

We acknowledge that experimental analyses of the final and intermediate products, as well as enzyme activity assays, are required to fully unravel the details of the metabolic pathways associated with a BGC. However, for the chosen classes of BGCs (NRPS, Type 1 PKS, and trans-AT PKS), we hypothesize that the information acquired from genome mining is sufficient to make *in silico* predictions that are biologically relevant. After assembling and evaluating the accuracy of the new pipeline presented in this work, we demonstrate its value towards high-throughput assessment of BGCs by reconstructing the metabolic pathways for 943 of the BGCs currently in MIBiG [34]. Furthermore, we predict the optimal single reaction inactivation (by gene knockout) strain-engineering strategy for natural product synthesis based on each BGC when introduced into a genome-scale metabolic model of *Streptomyces coelicolor*, a model organism among the *Actinobacteria* and a popular heterologous BGC expression host [15, 35].

## Results

We have developed the Biosynthetic Gene cluster Metabolic pathway Construction (BiGMeC) pipeline that leverages antiSMASH results to create the metabolic pathway corresponding to a PKS or NRPS biosynthetic gene cluster (Figure 1A). The pipeline details each enzymatic reaction of the metabolic pathway, including redox cofactors and energy demand. The results are stored in a format that is easily introduced into a GEM using popular tools for constraint-based reconstruction and analysis, such as cobrapy [36] or COBRA Toolbox [37].

**Figure 1.**
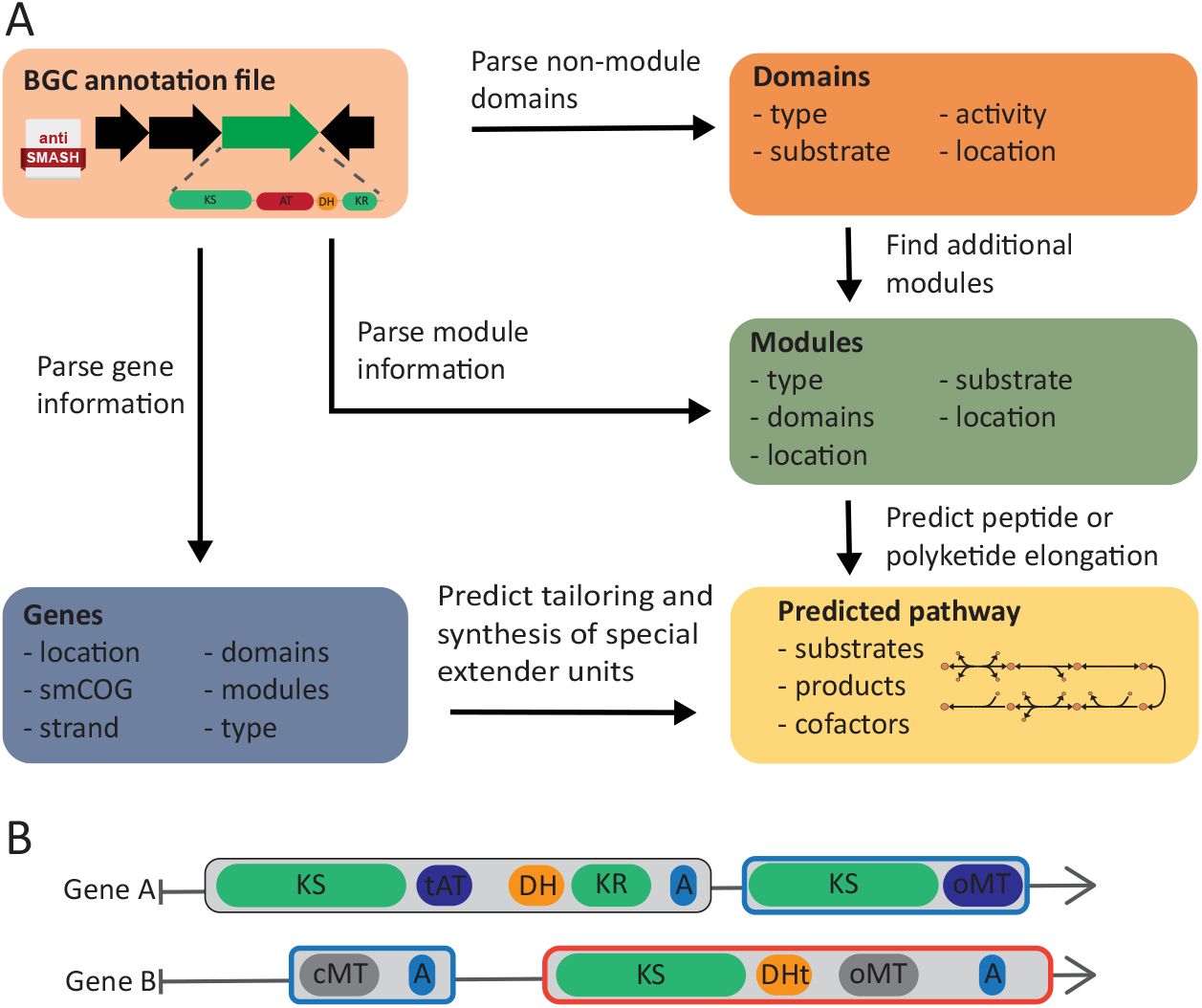
Overview of the BiGMeC pipeline. **A)** Schematic description of how the BGC annotation file produced by antiSMASH is parsed and used to construct the associated metabolic pathway. **B)** BiGMeC extends the rule-based identification of modules from antiSMASH with bridging modules and analysis of module activity, as exemplified here with this toy BGC: The last module in gene A and the first module in gene B (marked by green edges) constitute an active bridging module that is not identified by antiSMASH [12]. The last module on gene C (red edge color) are most often found to be inactive, a feature currently incorporated into BiGMeC.

The hallmarks of PKS- and NRPS-genes are adjacent functional domains that in total make up one or several modules that initiate, extend or cleave off the polyketide or peptide product, respectively [30, 33, 32]. The output from antiSMASH comprises information about these modules and their functional domains, and occasionally also the specific extender unit or chemical transformation associated with each functional domain [12]. The BiGMeC pipeline not only parses this information, but uses well-reasoned heuristics to handle deviations from canonical rules and cases where information is missing (see Materials and Methods). Improvements in determining module function includes identification of bridging modules in trans-AT PKSs and non-extending modules due to the presence of oMT domains [32] (Figure 1B).

We first assessed the accuracy of the BiGMeC pipeline by comparing its predictions with experimentally characterized and manually curated metabolic pathways. To this end, we compared the substrates, cofactors, and reaction products of each step of the metabolic pathway associated with eight well-characterized BGCs (Figure 2A, Supplemental Data 1). These BGCs cover a range of BGC classes, including Type 1 PKS, trans-AT PKS, NRPS and hybrids, and we believe they provide a test set that is sufficiently diverse to probe the pipeline for its strengths and weaknesses. Overall, BiGMeC appends the correct metabolic reaction for 72.8% (166/228) of the functional domains in all eight BGCs. Of these functional domains, BiGMeC chooses the correct extender unit for 81.3% (74/91) of the domains extending the peptide or polyketide. For all other domains, including chain initiation, reductive domains, methyltransferases and final tailoring reactions, the accuracy is 67.2% (92/137).

A large number of the incorrect predictions derive from wrong assignments of inactive KR domains by antiSMASH [12]. Across the eight closely inspected BGCs, KR domains are almost always active, but on several occasions antiSMASH predicts that these domains are inactive. Furthermore, this leads to incorrect assignment of the activity of succeeding DH and ER domains because they act on the functional moiety produced by the preceding domain. For the prediction of extender units, most incorrect assignments are due to the the inability of antiSMASH to determine if KS domains in trans-AT PKSs are active, e.g. for oocydin, only 10 of the apparent 16 KS domains are actually active. Another significant source of incorrect domains comes from the anabaenopeptin cluster that has two consecutive genes, each having two modules that initiate biosynthesis and perform first chain elongation, respectively, yielding two slightly different variants of the final compound. The BiGMeC pipeline treats these two genes as consecutive steps of the same pathway, and therefore, predicts too many chain elongations in the biosynthesis.

To investigate how much the errors in the constructed metabolic pathways affect model predictions, we introduced both the literature-based and the BiGMeC pathway reconstructions into the consensus GEM of *S. coelicolor* (Sco-GEM) [16] and compared the maximal production rate of the final compound (Figure 2B). In general, we observe quite similar rates for the eight BGCs (Pearson *ρ* = 0.75, *P* = 0.03), suggesting that the incorrect domains only have a minor impact on the predicted production rates. The offset in the production of leupyrrin likely comes from an incorrect starter unit while the offset in oocydin production is caused by a fairly large error in the predicted number of malonyl-CoA extender units (10 vs. 16).

**Figure 2.**
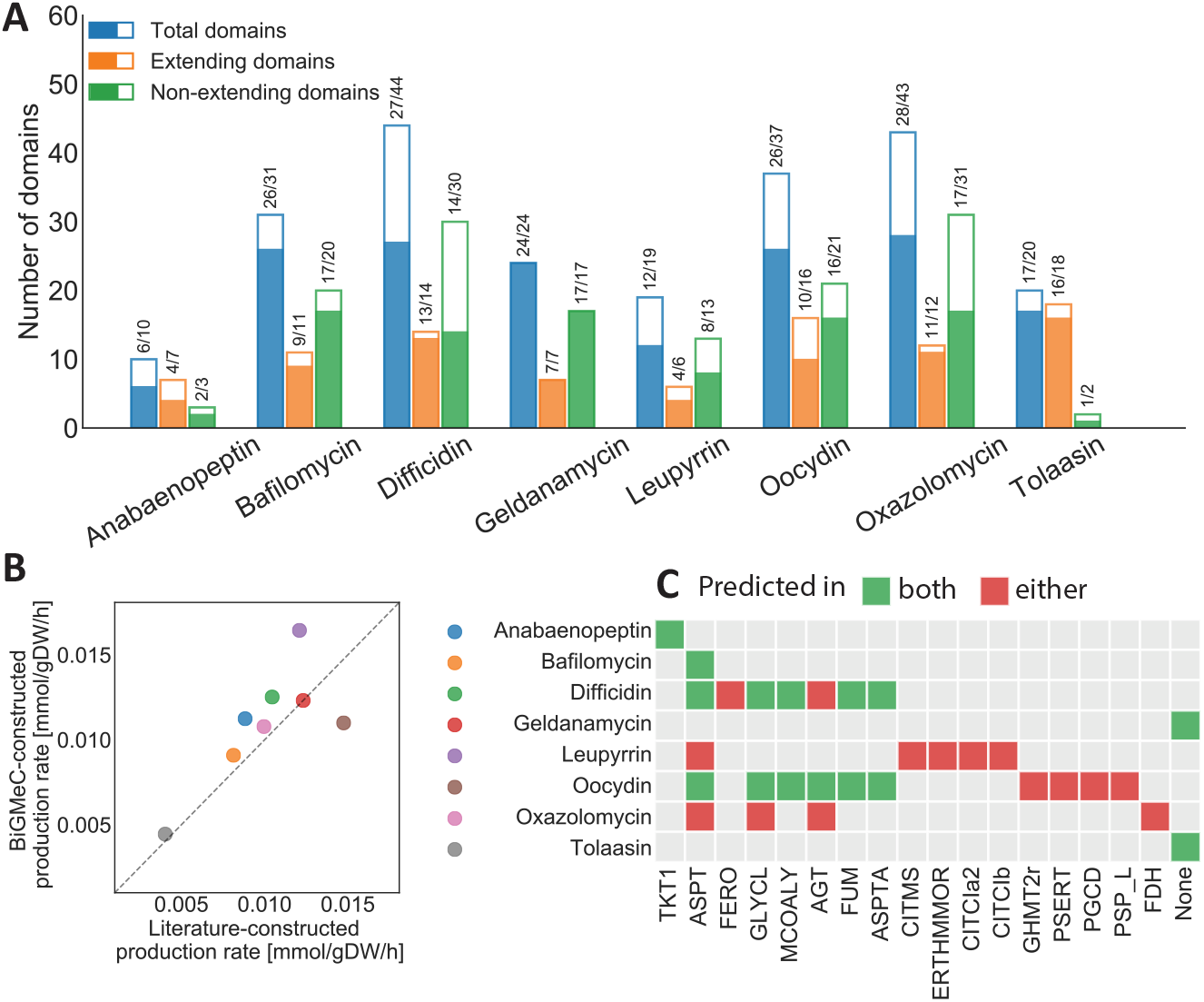
Analysis of BiGMeC prediction accuracy for eight selected BGCs. **A)** Barplot showing the number of correct domains when comparing BiGMeC-constructed pathways with pathways as they are detailed in the literature (Supplemental Data 1). The filled part of each bar, as well as the ratio printed above, shows the number of correct domains for each BGC. Extending domains comprises the domains that append an extender unit to the polyketide or peptide backbone, while the non-extending domains covers all other domains. **B)** Predicted maximum production rate when introduced into a *S. coelicolor* GEM. The x- and y-axis represent the maximal production rate using the metabolic pathway created based on literature or reconstructed with BiGMeC, respectively. **C)** This panel shows a comparison of the predicted reaction-knockout targets (x-axis) when using a metabolic pathway created based on literature or with BiGMeC. Similar predictions are shown as green tiles, while incorrect predictions (predicted in either but not both of the two cases) are shown as red tiles. The names of the model reaction IDs are: TKT1: transketolase; ASPT: aspartate ammonia-lyase; FERO: ferroxidase; GLYCL: Glycine Cleavage System; MCOALY: malyl-CoA lyase; AGT: alanine-glyoxylate aminotransferase; FUM: fumarase; ASPTA: aspartate transaminase; CITMS: (R)-citramalate synthase; ERTHMMOR: 3-isopropylmalate dehydrogenase; CITCIa2: (R)-2-Methylmalate hydro-lyase; CITCIb: 2-methylmaleate hydratase; GHMT2r: glycine hydroxymethyltransferase; PSERT: phosphoserine transaminase; PGCD: phosphoglycerate dehydrogenase; PSP_L: phosphoserine phosphatase; FDH: formate dehydrogenase.

The anticipated use of the developed pipeline towards strain engineering of expression hosts underscores the need to elucidate if model-based strain designs using BiGMeC-constructed pathways deviate from results using pathways reconstructed according to literature. To this end, we predicted optimal single-reaction knockout mutants that should increase the production rate of the associated product (Figure 2C). Note that, a reaction knockout is the practical implication of disrupting one or more of the genes encoding the enzyme catalyzing the corresponding reaction. For 6 out of 8 BGCs there is a good overlap between pairwise pathway reconstructions. This includes the cases of tolaasin and geldanamycin, where no knockout target is identified with either of the two pathway reconstructions.

To demonstrate the power of BiGMeC in high-throughput assessment of BGCs, we employed the pipeline on 1883 of the 1923 BGCs in the MIBiG database (version 2.0) [34]. For 40 of the 1923 BGCs, we could not obtain the antiSMASH output file because the link from MIBiG was broken. The 943 (50.1%) metabolic pathways that were successfully reconstructed with BiGMeC cover both fungi and a range of different bacteria (Figure 3A). Most clusters are either Type 1 PKS, NRPS, or hybrids of these two, and only 77 of the BGCs share similarity with trans-AT PKS (Figure 3B). The 940 remaining BGCs were not analyzed either because the BGC class was not covered by BiGMeC (such as RiPPs, terpenes, Type 2 and Type 3 PKSs) or because functional modules and domains were lacking in the results from antiSMASH.

**Figure 3.**
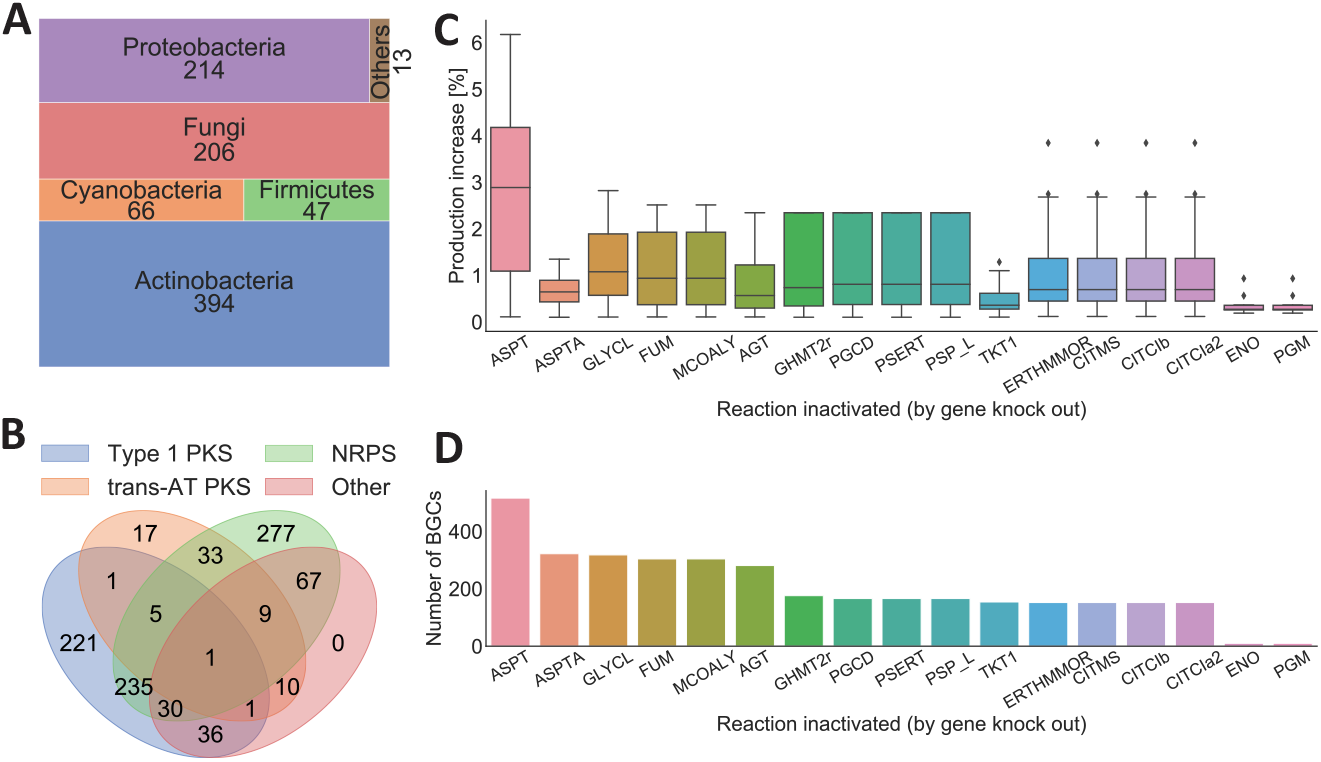
Automatic reconstruction and analysis of 943 BGCs from MIBiG. These BGCs cover a wide variety of hybrid BGCs and **B)** a range of different organisms. **C)** Box plot showing the increase in production for the 17 different reaction knockouts that increases the production of one or more of the analysed BGCs. **D)** Bar chart showing the number of BGCs where the knockout of each reaction is predicted to increase production of the target secondary metabolite. The 943 reconstructed pathways cover The names of the model reactions used in panel C and D: ASPT: aspartate ammonia-lyase; ASPTA: aspartate transaminase; GLYCL: Glycine Cleavage System; FUM: fumarase; MCOALY: malyl-CoA lyase; AGT: alanine-glyoxylate aminotransferase; GHMT2r: glycine hydroxymethyltransferase; PGCD: phosphoglycerate dehydrogenase; PSERT: phosphoserine transaminase; PSP_L: phosphoserine phosphatase; TKT1: transketolase; ERTHMMOR: 3-isopropylmalate dehydrogenase; CITMS: (R)-citramalate synthase; CITCIb: 2-methylmaleate hydratase; CITCIa2: (R)-2-Methylmalate hydro-lyase; ENO: enolase: PGM: phosphoglycerate mutase.

We introduced each of the 943 reconstructed pathways into Sco-GEM [16], and predicted single-reaction knockout strategies improving the production of the final pathway product. Surprisingly, only 17 different reactions were suggested as a knockout target in one or more of the 943 *in silico* heterologous expression experiments (Figure 3C and D). Of these 17 reactions, aspartate transaminase is predicted to provide on average the largest increase in production (Figure 3C) and is also the most frequently suggested candidate (Figure 3D). However, the predicted production increase is minor for all of the 17 suggested reactions, including aspartate transaminase, with a maximum increase of 6% relative to the wild-type production rate.

## Discussion

To make novel natural product pathways encoded by BGCs accessible to the constraint-based reconstruction and analysis framework, we have developed a pipeline that creates a draft reconstruction of the metabolic pathway encoded by a BGC. By applying the pipeline to 943 BGCs covering NRPSs, PKSs and NRPS-PKSs hybrids from a wide range of organisms we have demonstrated how the pipeline enables high-throughput assessment of potential candidates for heterologous expression. In an assessment of 943 BGCs, we explored general single-gene knockout strategies for increased heterologous production, and although we identify a set of 17 general targets, none provides a drastic increase in production. This result suggests that multiple knockouts, over-expression of genes, or strategies that perturb regulatory mechanisms are necessary to reroute a large amount of precursors from growth towards secondary metabolism, at least in the organism *S. coelicolor*. Although the accuracy of the BiGMeC pipeline is sufficient to make biologically relevant pathway reconstructions, this work has also revealed aspects where there are room for further improvement. Incorrect assignment of KS and KR domains as active or inactive is a large source of error in PKS metabolic pathways, and incorporation of the recently developed transATor algorithm would provide an improvement in this context [38]. Synthesis of rare precursors and tailoring of the polyketide or peptide succeeding the release from the multidomain enzyme complex are two other features with opportunity for improvement. Although the genes encoding enzymes responsible for the synthesis of rare precursors or for the post-release tailoring steps usually are contained in the BGC, neither their exact function nor their functional order can be accurately predicted. Therefore, the current pipeline relies in certain aspects on assumptions and heuristics that apply in general, but with several exceptions. However, with a continuous improvement in algorithms for annotation and identification of BGCs [12, 38, 39] and increased experimental characterization [34], current generalisations can develop into more accurate pathway reconstructions that encompass a larger range of deviations from canonical rules. Furthermore, as the knowledgebase and algorithms for annotation of iterative PKSs and ribosomally synthesised and post-translationally modified peptides improves, these types of BGCs represent obvious targets for further development. Further improvement should also aim to accept the output from other annotation software, such as PRISM [40], as well as expand to other metabolic pathways that are also encoded by gene clusters. An obvious candidate is exopolysaccharides, for which heterologous production can be an option for improved biotechnological production [41, 42].

## Conclusion

The BiGMeC pipeline is, to our knowledge, the first tool for automatic metabolic pathway reconstruction specifically targeting PKS and NRPS BGCs. Although the reconstructed pathways are not able to capture the entire diversity seen in the biosynthesis of NRPSs and PKSs [32, 30], the predicted production rates and reaction knockout targets are comparable to predictions provided using manually reconstructed pathways. Furthermore, the pipeline can aid model reconstruction efforts, both as a decent starting point for further manual curation and as a complement to standard model-reconstruction pipelines [43]. This is in particular relevant for organisms with a rich secondary metabolism, such as the *Actinobacteria* which are of utmost interest in drug discovery. We anticipate that the pipeline presented here can increase the use of GEMs in this context, e.g. to screen different combinations of BGCs and expression hosts or, as shown in this work, to explore strain-engineering opportunities. The pipeline is developed in an open source environment on GitHub and we encourage interested readers to engage in future development through pull request or by raising issues. We also encourage developers of genome mining tools and databases to converge towards standardized and consistent file formats, such as the Minimum information about a Biosynthetic Gene Cluster (MIBiG) initiative [34] This will ease the development and maintenance of downstream pipelines such as BiGMeC.

## Materials and Methods

### Software implementation

We developed the Biosynthetic Gene cluster to Metabolic pathway Constructer (BiGMeC) to translate information about PKS and NRPS BGCs to detailed outlines of the metabolic reactions governing the production of the associated secondary metabolites. The BiGMeC software and all other associated scripts are implemented in Python 3 and publicly available at https://github.com/AlmaasLab/BiGMeC. BiGMeC runs from a command-line interface and takes an annotated NRPS or PKS BGC in the format of a GenBank file as produced by antiSMASH 5.1 [12]. It leverages the included gene, domain, and module information to make a description of the enzymatic reactions encoded by the BGC, including substrate and co-factor usage (1A). BiGMeC uses a reference model as a library of metabolites and reactions, and in the current work, we have used Sco-GEM version 1.2.1, the consensus *S. coelicolor* GEM [16]. This model was obtained from https://github.com/SysBioChalmers/Sco-GEM.

The BiGMeC pipeline first parses information about the location and annotation of the genes and modules as annotated by antiSMASH from the GenBank file (Figure 1). If available, the gene information include strand, secondary metabolism Clusters of Orthologous Groups (smCOG) annotation [44], type of gene, extender unit, annotated functional domains and if the gene is a core gene or not. The core genes synthesize the core structure of the PKS or NRPS molecule. The module information contains details about the type of module and its functional domains. Then, the pipeline assesses the presence and order of domains not included in a module, e.g. special load or bridging modules (in trans-AT PKS, Figure 1B) [32], and combines these domains into functional modules when possible. The peptide or polyketide backbone is subsequently constructed based on the order of the identified domains and the function of each domain within each module. The reactions associated with the functional domains are listed in Table 1. Domains in the BGC that are not contained in a module are assumed to not affect the backbone structure. If a terminating domain (thioesterase or thioester reductase) domain is encountered, no further chain elongations are carried out. The activity of reducing domains (DH, ER, KR) are based on the annotation of the KR domain from antiSMASH. Tailoring reactions post PKS synthesis are predicted from the smCOG annotations of each gene. The currently implemented tailoring reactions relate to the smCOGs 1256, 1084, 1002, 1109 and 1062 and includes glycosylation, glycosyltransferase and incorporation of 2-Amino-3-hydroxycyclopent-2-enone (Supplemental Material 1).

**Table 1.**
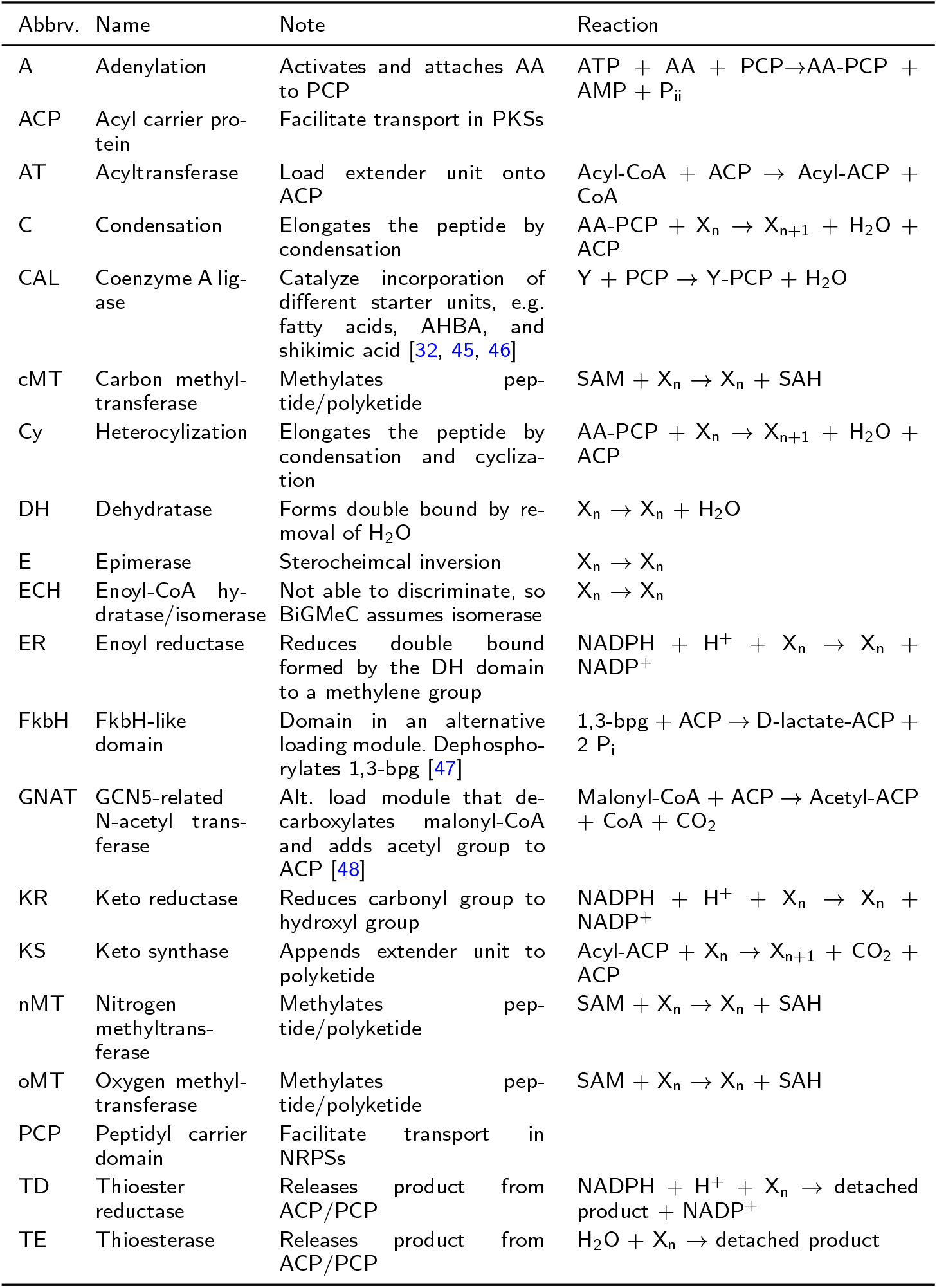
List of domains and associated reactions as implemented in BiGMeC. The peptide or polyketide backbone is referred to as Xn, and in reactions that extend the backbone we refer to the elongated backbone as X_n+1_. Other abbreviations used: AA: generic amono acid; 1,3-bpg: 1,3 biphosphoglycerate; CoA: Coenzyme A; P_ii_: diphosphate; SAH: S-Adenosyl-L-homocysteine; SAM: S-Adenosyl methionine; Y: generic starter unit.

| Abbrev. | Name | Note | Reaction |
| --- | --- | --- | --- |
| A | Adenylation | Activates and attaches AA to PCP | $ATP + AA + PCP \rightarrow AA-PCP + AMP + P_{ii}$ |
| ACP | Acyl carrier protein | Facilitate transport in PKSs |  |
| AT | Acyltransferase | Load extender unit onto ACP | $Acyl-CoA + ACP \rightarrow Acyl-ACP + CoA$ |
| C | Condensation | Elongates the peptide by condensation | $AA-PCP + X_n \rightarrow X_{n+1} + H_2O + ACP$ |
| CAL | Coenzyme A ligase | Catalyze incorporation of different starter units, e.g. fatty acids, AHBA, and shikimic acid [32, 45, 46] | $Y + PCP \rightarrow Y-PCP + H_2O$ |
| cMT | Carbon methyltransferase | Methylates peptide/polyketide | $SAM + X_n \rightarrow X_n + SAH$ |
| Cy | Heterocyclization | Elongates the peptide by condensation and cyclization | $AA-PCP + X_n \rightarrow X_{n+1} + H_2O + ACP$ |
| DH | Dehydratase | Forms double bond by removal of $H_2O$ | $X_n \rightarrow X_n + H_2O$ |
| E | Epimerase | Stereochemical inversion | $X_n \rightarrow X_n$ |
| ECH | Enoyl-CoA hydratase/isomerase | Not able to discriminate, so BiGMeC assumes isomerase | $X_n \rightarrow X_n$ |
| ER | Enoyl reductase | Reduces double bond formed by the DH domain to a methylene group | $NADPH + H^+ + X_n \rightarrow X_n + NADP^+$ |
| FkbH | FkbH-like domain | Domain in an alternative loading module. Dephosphorylates 1,3-bpg [47] | $1,3-bpg + ACP \rightarrow D-lactate-ACP + 2 P_i$ |
| GNAT | GCN5-related N-acetyl transferase | Alt. load module that decarboxylates malonyl-CoA and adds acetyl group to ACP [48] | $Malonyl-CoA + ACP \rightarrow Acetyl-ACP + CoA + CO_2$ |
| KR | Keto reductase | Reduces carbonyl group to hydroxyl group | $NADPH + H^+ + X_n \rightarrow X_n + NADP^+$ |
| KS | Keto synthase | Appends extender unit to polyketide | $Acyl-ACP + X_n \rightarrow X_{n+1} + CO_2 + ACP$ |
| nMT | Nitrogen methyltransferase | Methylates peptide/polyketide | $SAM + X_n \rightarrow X_n + SAH$ |
| oMT | Oxygen methyltransferase | Methylates peptide/polyketide | $SAM + X_n \rightarrow X_n + SAH$ |
| PCP | Peptidyl carrier domain | Facilitate transport in NRPSs |  |
| TD | Thioester reductase | Releases product from ACP/PCP | $NADPH + H^+ + X_n \rightarrow detached product + NADP^+$ |
| TE | Thioesterase | Releases product from ACP/PCP | $H_2O + X_n \rightarrow detached product$ |

Rare extender units appear in both PKS and NRPS biosynthesis. The synthesis of rare extender units are usually carried out by genes in the BGC [49], and we therefore include the synthesis of the most common rare extender units (not in the reference library) when necessary. This includes hydroxyphenylglycine, beta-hydroxytyrosine, 2-aminobutyric acid, pipecolic acid, dihydroxyphenylglycine and 3-amino-5-hydroxybenzoate[45]. Synthesis of the rare extender unit methoxymalonyl-ACP [49] is based on the presence of genes with specific smCOG annotations (Supplemental Material 1). For the remaining rare extender units, or in the case of missing information or nonspecific antiSMASH annotation, we use a conservative approach where a generic amino acid is used as the extender unit in NRPS modules and malonyl-CoA is used in PKS modules. In the case of using a generic amino acid as the extender unit, we add a set of pseudo-reactions that can convert every proteogenic amino acid into this generic molecule to ensure that the biosynthethic pathway is functional.

The pipeline also handles a number of deviations from the canonical rules, for example the deactivation of the KS domain often seen in modules containing Omethyltransferases [32]. Furthermore, it is found that the presence of a C domain in the initiating NRPS module acylates the initial amino acid [50, 31]. Both in tolaasin [51] and surfactin, currently the best studied example of this type of NRPS initiation, the acylating agent is a CoA-activated *β*-hydroxy fatty acid [50, 52]. It is likely that the C-domain has a strong selectivity for a specific acylating agent, but since this specificity is not identified by antiSMASH we use a generic fatty acid molecule. A third example of exception thatis handled are bridging modules in trans-AT PKSs where the KS domain is encoded in the first gene and a DH and ACP domain follows immediately on the second gene. These modules are called dehydratase docking domains (DHD) and are usually not active [32].

### Evaluation of BiCMeC pipeline

To evaluate how well biosynthetic pathways can be constructed solely based on antiSMASH data we compared BiGMeC-constructed pathways with literature-based reconstructions for 8 different BGCs, covering different species and classes of BGCs (Supplemental Data 1). The 8 BGCs were (MIBiG ID in parenthesis): bafilomycin from *Streptomyces lohi* [53, 54, 55] (BGC0000028), geldanamycin from *Strepto-myces hygroscopicus* [56, 57, 58] (BGC0000066), difficidin from *Bacillus velezensis FZB42* [59, 60] (BGC0000176), oocydin from *Serratia plymuthica* [32, 61] (BGC0001032), oxazolomycin from *Streptomyces albus* [60, 62] (BGC0001106), leupyrrin from *Sorangium cellulosum* [63] (BGC0000380), anabaenopeptin from *Anabaena sp. 90* [64] (BGC0000302) and tolaasin from *Pseudomonas costantinii* [51] (BGC0000447). For each domain in each of the 8 different BGCs we compared the BiGMeC-constructed reaction with the *real* reaction, i.e. the associated reaction as described in the literature. When clearly defined in the literature, tailoring reactions were included, but we focused on the synthesis of the core peptide/polyketide. The very complex tailoring of leupyrrin [63] was not included.

An initial evaluation was performed by counting the number of correct domains (Figure 2A). The total number of domains include all domains either predicted by BiGMeC or described in the literature, and the correct predictions include both true positives and true negatives. Next, we incorporated the BiGMeC and literature-based pathway reconstructions into Sco-GEM and predicted the maximum production rate of the secondary metabolite produced by each pathway (Figure 2B). To do so, we performed Flux Balance Analysis (FBA) [65, 66] in cobrapy [36] with the final reaction of the BGC encoded pathway as objective and with growth limited to minimum 90% of the maximum value. The growth and production was simulated in a growth medium with glucose and ammonium as the sole carbon and nitrogen source, respectively, and with a maximum glucose uptake rate of 0.8 mmol gDW^−1^ h^−1^. We did not constrain the uptake of ammonium, sulphate, phosphate, oxygen and metal ions. Finally, using both the BiCMeC and literature-based pathway reconstructions, we predicted reaction inactivation targets (by gene knockout) that would increase the production of the associated compound, with a maximum growth rate reduction of 50% (Figure 2C). We limited the set of possible reaction targets to non-essential gene-annotated reactions. The search for optimal knockouts was carried out in a brute-force manner: we conducted an iterative knockout of each reaction (within the predefined set of possible reactions) and, first used FBA to predict the maximum growth of the mutant phenotype, and secondly predict the maximum production rate at 99.9% of the knockout-mutant’s maximum growth rate. All knockouts that resulted in more than 0.1% increase in production rate compared to the wild-type were considered knockout candidates.

### Large-scale reconstruction of BGC pathways

To demonstrate the value and efficiency enabled by BiGMeC we applied this pipeline to all relevant BGCs from the MIBiG database [34]. To get the antiSMASH-generated output for all BGCs in MIBiG we automatically downloaded all GenBank-files with a url on the form: https://mibig.secondarymetabolites.org/repository/BGC0000001/generated/BGC0000001.1.region001.gbk, with the MIBiG ID ranging from BGC0000001 to BGC0002057. The MIBiG database currently reports on a total of 1923 BGCs but due to different reasons (e.g. missing entries) we could only obtain the antiSMASH result for 1883 of the entries. For all BGCs at least annotated to either Type 1 PKS, trans-AT PKS or NRPS we used the BiGMeC pipeline to reconstruct the corresponding metabolic pathway. We predicted optimal knockout strategies for each of successfully constructed pathway using the same procedure as described for the 8 BGCs used to evaluate the BigMeC pipeline.

## Declarations

Ethics approval and consent to participate Not applicable.

## Consent for publication

Not applicable.

## Availability of data and materials

The BiGMeC pipeline and the data analysed/generated during the current study is available at https://github.com/AlmaasLab/BiGMeC. We have also deposited the latest version of the repository to Zenodo (https://doi.org/10.5281/zenodo.4287511) to ensure persistent access.

## Author’s contributions

Conceptualization, SS, EA; Methodology and Software, SS, FF; Validation and Formal Analysis, SS, FF; Data curation SS, FF; Writing - Original Draft, SS, FF; Reviewing and editing, SS, EA, FF, AW; Visualization SS; Supervision SS, EA; Project Administration, AW, EA; Funding Acquisition, AW, EA.

## Acknowledgements

Not applicable.

## Additional Files

### Supplemental Data 1

Detailed comparison of 8 BGCs for evaluation the accuracy of the BiGMeC pipeline.

### Supplemental Material 1

Details on tailoring reactions and synthesis of methoxymalonyl-ACP, and description of the analysis used to make heuristics that indicate the presence of these reactions from smCOG annotations.

