## Supplemental Material 1 for "Automatic reconstruction of metabolic pathways from identified biosynthetic gene clusters"

### Tailoring reactions

We surveyed gene clusters in MIBiG [1] to explore how specific smCOG gene annotations could be used to determine the tailoring reactions associated with the biosynthesis of a polyketide or non-ribosomal peptide synthetase: we compared the presence of specific secondary metabolism Clusters of Orthologous Groups (smCOGs) [2] annotations with the presence of different tailoring reactions as described in the literature. This led to three fairly robust heuristic that were implemented into the BiGMeC pipeline (Table S1). The data that these heuristics are based on are further described in separate sections below.

Table S1: Boolean logic used to determine tailoring reactions from smCOG gene annotation. The peptide or polyketide that is modified by the tailoring reactions is denoted by $X_n$, both prior and subsequent to the tailoring. Other abbreviations used in the table: bpg: 1,3-biphopshoglycerate; $P_i$: phosphate; $P_{ii}$: diphosphate; CoA: Coenzyme A.

| smCOG logic | Reaction |
|---|---|
| 1256 and 1084 | bpg + NADH + $H^+$ + $X_n$ → $X_n$ + 2 $P_i$ + $NAD^+$ |
| 1002 and 1109 | glycine + succinyl-CoA + ATP + $X_n$ → $X_n$ + AMP + $P_{ii}$ + $CO_2$ + $H_2O$ + CoA |
| 1062 | glucose 6-phophate + $X_n$ → $X_n$ + $P_i$ + $H^+$ |

#### Addition of glycerate

Tailoring by addition of glycerate from 1,3-biphopshoglycerate was identified from the presence of the smCOGs 1256 (FkbH like protein) and 1084 (3-oxoacyl-(acyl carrier protein) synthase III). Across all BGCs in MIBiG, this heuristic provides a correct tailoring reaction in 10 out of 11 cases (Table S2). The single error derives from BGC0000082 which still incorporates a glycerate unit, but lacking the smCOG 1084 annotation.

#### Incorporation of 2-Amino-3-hydroxycyclopent-2-enone

The tailoring reaction that incorporate 2-Amino-3-hydroxycyclopent-2-enone synthesized from succinyl-CoA and glycine [12], was identified from the

Table S2: Listing of BGCs in MIBiG used to determine when the addition of glycerate is used to tailor the polyketide or peptide. These are all BGCs in MIBiG containing the FkbH domain and adds glycerate. Of these 11 BGCs, 10 are annotated with the smCOG 1084.

| MIBiG ID | Product | 1084 present | Reference |
|---|---|---|---|
| BGC0000001 | Abyssomycin | Yes | [3] |
| BGC0000036 | Chlorothricin | Yes | [4] |
| BGC0000082 | Kijanimicin | No | [5] |
| BGC0000133 | Quartromicin | Yes | [6] |
| BGC0000140 | 4H3H2HMe2HF5 | Yes | [6] |
| BGC0000162 | Tetrocarcin | Yes | [7] |
| BGC0000164 | Tetronomycin | Yes | [8] |
| BGC0001004 | Lobophorin B | Yes | [9] |
| BGC0001183 | Lobophorin A | Yes | [9] |
| BGC0001204 | Versipelostatin | Yes | [10] |
| BGC0001288 | Maklamicin | Yes | [11] |

presence of the smCOGs of 1109 (8-amino-7-oxononanoate synthase) and a neighboring 1002 (AMP-dependent ligase and synthetase). This pair of smCOG annotations were present in 8 BGCs in MIBiG, of which 5 features a tailoring reaction that incorporates 2-Amino-3-hydroxycyclopent-2-enone (Table S3). For one of the BGCs (BGC0001420) the available literature was not sufficient to make a clear decision about the compound tailoring. The remaining two BGCs (BGC0000091 and BGC0001063) feature a similar tailoring reaction where the succinyl-CoA precursor is replaced by several malonyl-CoA units [13, 14].

### Glycosylation

Glycosylation of the peptide or polyketide is identified by the presence of smCOG 1062 (glycosyltransferase), and as one might expect the number of incorporated sugar monomers increase with increasing number of smCOG 1062 annotations (Table S4). Based on 40 BGC synthesis pathways that incorporate sugar monomers (we consider Komodoquinone B as an outlier), we find that the correlation between the number of incorporated sugar monomers and the number of smCOG 1062 annotations is 0.78 (p value = 3e-9). This demonstrate that most, but not all, of the glycosyltransferases are active. Based on this result the BiGMeC pipeline therefore assumes that all glycosyltransferases are active, i.e. a 1 to 1 relationship between the number of smCOG 1062 and glycosyltransferase tailoring reactions.

Table S3: Listing of BGCs in MIBiG used to determine when the peptide or polyketide is tailored by the incorporation of 2-Amino-3-hydroxycyclopent-2-enone synthesized from succinyl-CoA and glycine [12]. All of these BGCs are annotated with the smCOG 1002 and 1109 on adjacent genes. The "Correct" column indicate if these synthesis of the molecules associated with each BGC incorporates 2-Amino-3-hydroxycyclopent-2-enone.

| MIBiG ID | Product | Correct | Reference |
|---|---|---|---|
| BGC0000028 | Bafilomycin | Yes | [15] |
| BGC0000091 | Marineosin | No | [13] |
| BGC0000187 | Asukamycin | Yes | [16] |
| BGC0000213 | Colabomycin | Yes | [17] |
| BGC0001063 | Undecylprodigiosin | No | [14] |
| BGC0001298 | Annimycin | Yes | [18] |
| BGC0001420 | Myxochromide | Unknown | [19] |
| BGC0001740 | Phthoxazolin | Yes | [20] |

Table S4: Listing of BGCs in MIBiG that incorporates sugar monomers by glycosyltransferase in one of the tailoring steps. The "# in BGC" and "# in pathway" columns display the number of smCOG 1062 annotations the BGC and the number of active glycosyltransferases in the synthesis of the associated compound.

| MIBiG ID | Product | # in BGC | # in pathway | Reference |
|---|---|---|---|---|
| BGC0000002 | Aculeximycin | 5 | 8 | [21] |
| BGC0000021 | Apoptolidin | 3 | 2 | [22] |
| BGC0000033 | Calicheamicin | 4 | 4 | [23] |
| BGC0000034 | Candicidin | 1 | 1 | [24] |
| BGC0000035 | Chalcomycin | 2 | 1 | [25] |
| BGC0000036 | Chlorothricin | 2 | 2 | [4] |
| BGC0000042 | Cremimycin | 1 | 1 | [26] |
| BGC0000052 | ECO-02301 | 1 | 1 | [27] |
| BGC0000054 | Erythromycin B | 2 | 2 | [28] |
| BGC0000078 | Inecidine | 2 | 3 | [29] |
| BGC0000081 | Kedarcidin | 2 | 2 | [30] |
| BGC0000082 | Kijanimicin | 5 | 5 | [5] |
| BGC0000085 | Lankamycin | 2 | 3 | [31] |
| BGC0000092 | Megalomicins | 3 | 3 | [32] |
| BGC0000096 | Midecamicin | 2 | 1 | [33] |
| BGC0000102 | Mycinamicin II | 2 | 1 | [34] |
| BGC0000105 | Nanchangmycin | 1 | 1 | [35] |
| BGC0000108 | Natamycin | 1 | 1 | [36] |
| BGC0000115 | Nystatin A1 | 1 | 1 | [37] |
| BGC0000136 | Rifamycin | 1 | 1 | [38] |
| BGC0000141 | Rubradirin | 1 | 2 | [39] |
| BGC0000148 | A83543A | 2 | 2 | [40] |
| BGC0000151 | Stambomycin A | 1 | 1 | [41] |
| BGC0000162 | Tetrocarcin A | 5 | 4 | [7] |
| BGC0000165 | Tiacumicin B | 2 | 2 | [42] |
| BGC0000167 | Vicenistatin | 1 | 1 | [43] |
| BGC0000197 | Aranciamycin | 1 | 1 | [44] |
| BGC0000198 | Arenimycin A/B/C | 2 | 2 | [45] |
| BGC0000199 | ArimetamycinA | 2 | 3 | [46] |
| BGC0000199 | ArimetamycinB | 1 | 3 | [46] |
| BGC0000199 | ArimetamycinC | 1 | 3 | [46] |
| BGC0000200 | Arixanthomycin A | 1 | 2 | [47] |
| BGC0000203 | BE-7585A | 3 | 1 | [48] |
| BGC0000208 | Chelocardin | 0 | 1 | [49] |
| BGC0000210 | Chromomycin A3 | 5 | 4 | [50] |
| BGC0001183 | Lobophorin | 3 | 4 | [9] |
| BGC0001452 | Sipanmycin | 2 | 4 | [51] |
| BGC0001522 | Auroramycin | 2 | 4 | [52] |
| BGC0001619 | Ibomycin | 6 | 7 | [53] |
| BGC0001851 | Komodoquinone B | 0 | 5 | [54] |
| BGC0002033 | Spiramycin | 3 | 4 | [55] |

### Rare extender units

The smCOG gene annotations were also leveraged to determine if the genes encoding enzymes responsible for the synthesis of the rare polyketide precursor $P_{ii}$malonyl-CoA were present in the BGC. This synthesis pathway was identified by the presence of smCOG 1256 (FkbH like protein) and smCOG 1095 (3-hydroxybutyryl-CoA dehydrogenase), see Table S5. This heuristic predicts the correct extender unit in 16 of the 24 relevant BGCs. The relevant BGC are selected from the MIBiG database based those that contains a gene with the smCOG 1245 and incorporates one or more rare extender unit. Seven of the 8 incorrect predictions are due to BGCS incorporating hydroxymalonyl-ACP and not methoxymalonyl-ACP, and the last error derives from BGC0000090 which incorporates methoxymalonyl-ACP, but lacks the smcOG 1095 annotation. Because we are currently not able to discriminate the synthesis of hydroxymalonyl-CoA from the synthesis of methoxymalonyl-ACP based on smCOG annoations, the BiGMeC pipeline assumes that the rare extender unit is methoxymalonyl-ACP. While this determines the incorporation of the reactions synthesising methoxymalonyl-ACP [56], the incorporation of methoxymalonyl-ACP as an extender unit only occurs if this requirement is fulfilled and methoxymalonyl-ACP is the extender unit as suggested by antiSMASH [57].

Table S5: List of BGCs in MIBiG that contains a gene with the smCOG 1245 annotation and incorporates a rare extender unit.

| MIBiG ID | Product | Substrate | 1095 present | Reference |
|---|---|---|---|---|
| BGC0000020 | Actinosynnema | Hydroxymalonyl-ACP | Yes | [58] |
| BGC0000021 | Apoptolidin | Methoxymalonyl-ACP | Yes | [22] |
| BGC0000028 | Bafilomycin | Methoxymalonyl-ACP | Yes | [15] |
| BGC0000040 | Concanamycin A | Methoxymalonyl-ACP | Yes | [59] |
| BGC0000065 | Rustmicin | Methoxymalonyl-ACP | Yes | [60] |
| BGC0000066 | Geldanamycin | Methoxymalonyl-ACP | Yes | [61] |
| BGC0000074 | Herbimycin A | Methoxymalonyl-ACP | Yes | [38] |
| BGC0000078 | Incednine | Methoxymalonyl-ACP | Yes | [29] |
| BGC0000090 | Macbecin | Methoxymalonyl-ACP | No | [62] |
| BGC0000096 | Midecamycin | Methoxymalonyl-ACP | Yes | [33] |
| BGC0000159 | Tautomycin | Methoxymalonyl-ACP | Yes | [63] |
| BGC0000970 | Chondrochloren A | Methoxymalonyl-ACP | Yes | [64] |
| BGC0001034 | Pellasoren | Methoxymalonyl-ACP | Yes | [65] |
| BGC0001054 | Xenocoumacin | Hydroxymalonyl-ACP | Yes | [66] |
| BGC0001059 | Zwittermycin A | Hydroxymalonyl-ACP | Yes | [67] |
| BGC0001106 | Oxazolomycin B | Methoxymalonyl-ACP | Yes | [68] |
| BGC0001348 | JBIR-100 | Methoxymalonyl-ACP | Yes | [69] |
| BGC0001511 | Ansamitocin P-3 | Methoxymalonyl-ACP | Yes | [70] |
| BGC0001537 | Butyrolactol A | Hydroxymalonyl-ACP | Yes | [71] |
| BGC0001902 | Bengamide | Hydroxymalonyl-ACP | Yes | [72] |
| BGC0001956 | Miharamycin A | Hydroxymalonyl-ACP | Yes | [73] |
| BGC0001957 | Amipurimycin | Hydroxymalonyl-ACP | Yes | [73] |
| BGC0002011 | Ansacarbamitocin A | Methoxymalonyl-ACP | Yes | [74] |
| BGC0002033 | Spiramycin | Methoxymalonyl-ACP | Yes | [55] |
